# An Engineered Human-Antibody Fragment with Fentanyl Pan-Specificity that Reverses Carfentanil-Induced Respiratory Depression

**DOI:** 10.1101/2023.07.04.547721

**Authors:** Lisa M. Eubanks, Tossapol Pholcharee, David Oyen, Yoshihiro Natori, Bin Zhou, Ian A. Wilson, Kim D. Janda

## Abstract

The opioid overdose crisis primarily driven by potent synthetic opioids resulted in more than 500,000 deaths in the US over the last 20 years. Though naloxone, a short acting medication, remains the primary treatment option for temporarily reversing opioid overdose effects, alternative countermeasures are needed. Monoclonal antibodies present a versatile therapeutic opportunity that can be tailored for synthetic opioids and that can help prevent post-treatment renarcotization. The ultrapotent analog carfentanil, is especially concerning due to its unique pharmacological properties. With this in mind, we generated a fully human antibody through a drug-specific B cell sorting strategy with a combination of carfentanil and fentanyl probes. The resulting pan-specific antibody was further optimized through scFv phage display. This antibody, C10-S66K, displays high affinity to carfentanil, fentanyl, and other analogs, and reversed carfentanil-induced respiratory depression. Additionally, x-ray crystal structures with carfentanil and fentanyl bound provided structural insight into key drug:antibody interactions.

---

The opioid epidemic has been characterized as having three distinct but overlapping waves with the first wave starting in 1999 with increased prescribing of opioid medications for the treatment of pain. Overdose deaths involving pharmaceutical opioids sharply increased from 3,442 in 1999 to a high of 17,029 in 2017, as a direct result of over-prescription of opioids backed by the notion that risk of addiction was very low.1 The second wave marked by an increase in deaths involving heroin emerging around 2010 was propelled by the tightening of regulations concerning opioid medications and the acknowledgement of addiction liabilities. Heroin overdose death rates nearly quadrupled between 2002 and 2013 and continued to rise, peaking at 15,482 in 2017.^1^ Although overdose deaths connected to opioid prescriptions and heroin have declined in recent years, overall opioid-related deaths still topped 80,000 in the US in 2021. Synthetic opioids are responsible for the third wave beginning in 2013 and have accounted for the vast majority of opioid-related deaths over the last 5 years with as much as 82% in 2020.^1^

Synthetic opioids have traditionally been used in medical settings as analgesics, sedatives and for pain management, but recently have flooded the illegal drug market due to their extreme potencies and cheap production costs. Fentanyl and related synthetic opioids, the main culprits, are opioid receptor agonists that selectively bind μ-opioid receptors (MORs) in the brain. Activation of these MORs produce analgesia, sedation, miosis, euphoria and respiratory depression, the latter being the primary cause of overdose and death.^2-3^

Fentanyl and its structurally related analogs comprise the 4-anilidopiperidine class of synthetic opioids, a class of compounds with relatively high lipophilicity and low molecular weight that can readily cross the blood brain barrier (BBB).^4^ Compared to morphine and heroin, fentanyl is approximately 100-fold and 10-fold more potent, respectively (Figure 1). Among this class of synthetics, MOR binding affinity generally correlates with drug potency and minor changes in structure have resulted in major shifts in activity. Simple addition of a methyl ester to fentanyl’s piperidine ring results in a 100-fold increase in potency for carfentanil, the most potent of the analogs detected in the US (Figure 1).^5-6^ This drastic change in strength is reflected in the MOR binding affinities for fentanyl (K_µ_= 1.2 nM) and carfentanil (K_µ_= 0.024 nM) that in part account for their pharmacological differences.^7^

**Figure 1.**
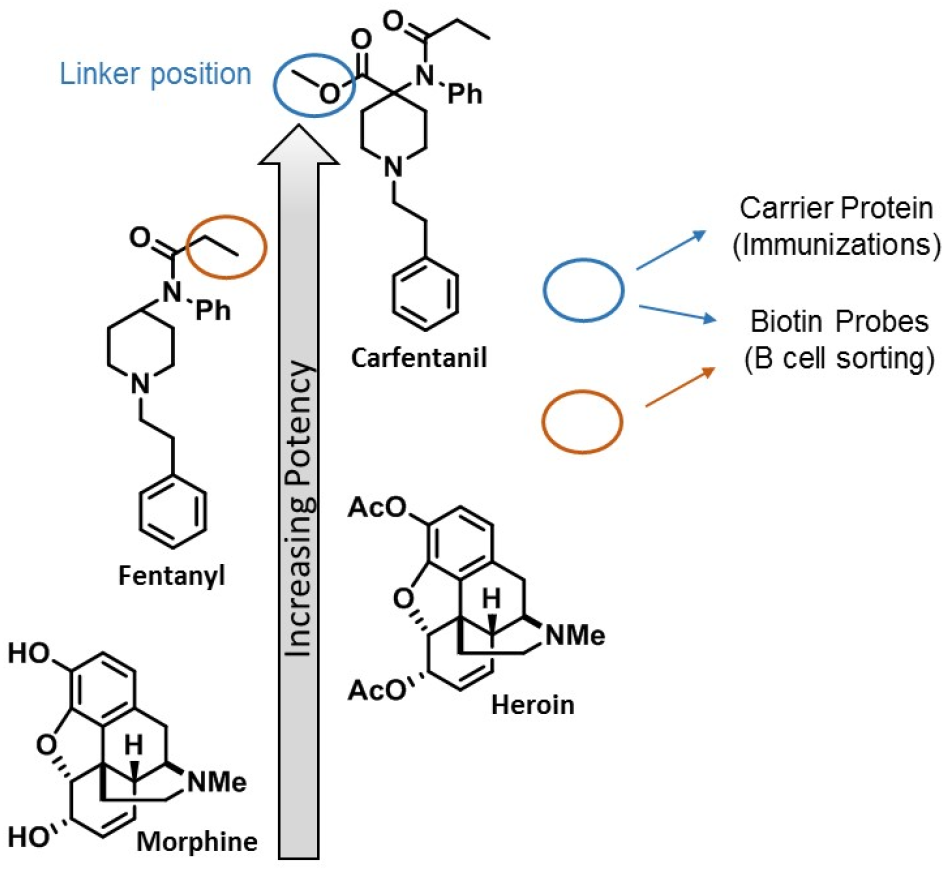
Chemical structures of opioids and respective potencies. Linker positions of the carfentanil immunoconjugate and drug-specific B cell sorting probes used for monoclonal antibody production and selection are indicated.

Fentanyl and carfentanil have both been added to counterfeit pills, heroin, cocaine and methamphetamine by drug dealers to increase profits. Unfortunately, most users are unaware that the illicit drugs they are acquiring contain one or more powerful adulterants. This new trend has proven deadly, resulting in skyrocketing overdose deaths attributed to opioid contaminants. In fact, the Food and Drug Administration (FDA) decided to pull carfentanil off the market in an attempt to curtail access to this ultrapotent opioid after it was routinely found in heroin supplies. Carfentanil is the second most frequently found opioid adulterant next to fentanyl and has recently been reported throughout the US as well as Europe and Canada.^8^ Carfentanil is also listed as a chemical threat agent by the Countermeasures Against Chemical Threats (CounterAct) Program, an NIH-initiative that supports the development of therapeutics against chemical warfare agents. If weaponized, carfentanil has the potential to cause mass casualties due to its tremendous toxicity.^9^

Treatment of carfentanil overdose presents a unique set of innate challenges due to extreme potency, high receptor affinity, long half-life (t_1/2_), slow drug redistribution, and profound deepening of respiratory depression. Naloxone, a MOR antagonist, is the only Food and Drug Administration (FDA)-approved medication to treat opioid overdose but may not effectively reverse carfentanil-induced intoxication. The short-lasting activity of naloxone requires much higher and frequent doses to maintain detoxification and prevent renarconization, a rapid return to a drug overdose state. In light of naloxone’s limitations, alternative countermeasures such as monoclonal antibodies (mAb) are currently being developed that can decrease the drug at the site of action.^10^

Murine and chimeric mAbs have been described previously against members of the opioid family including morphine, heroin, heroin’s metabolite 6-acetylmorphine and fentanyl.^11-15^ However, from a therapeutic standpoint, the presence of non-human derived antibody regions is a liability often triggering an adverse immune response. Therefore, we began a research program centered on carfentanil with the goal of generating fully human antibodies that can be used to treat drug overdose and that are cross-reactive with the prevalent fentanyl and its analogs. The general strategy for this endeavor consisted of immunization with an appropriate drug immunoconjugate, sorting antigen-specific single B cells, cloning and expressing mAbs of interest, and kinetic characterization of positive mAb clones (Figure 2).

**Figure 2.**
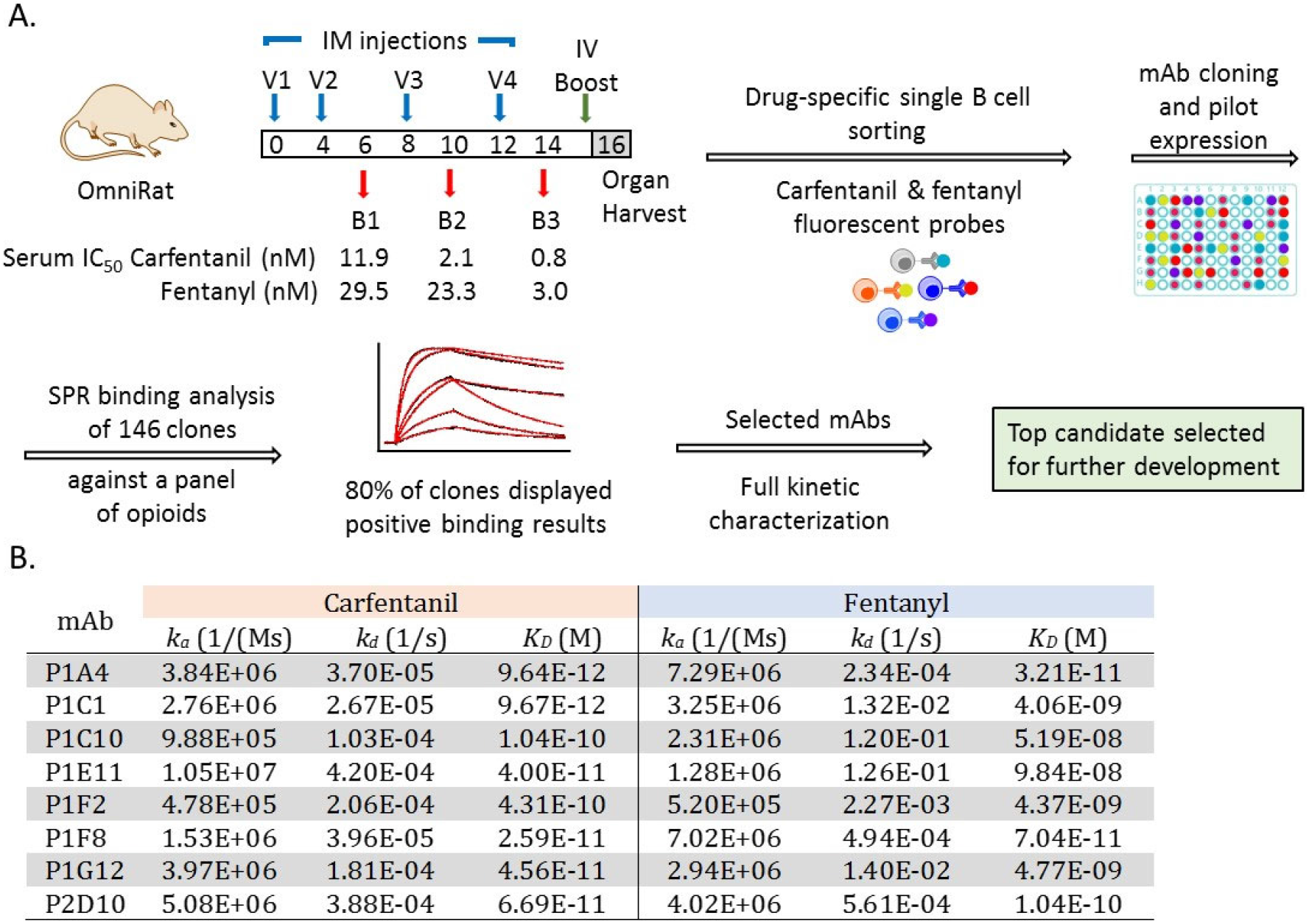
General strategy for generation and selection of monoclonal antibodies (mAb) against carfentanil, fentanyl and related synthetic opioids. (A) Overview of the process leading to the selection of our lead anti-opioid mAb including immunization of IgG-transgenic rats, single B cell sorting and binding analysis of single antibody clones. (B) Kinetic rate constants for lead mAbs binding to carfentanil and fentanyl measured by surface plasmon resonance

To this end, we exploited human IgG-transgenic rats that express human antibody recognition domains and facilitate the production of fully human antibodies of optimal affinity and diversity.^16-17^ Rats were immunized with our carfentanil hapten, Carfen-*ester*, that has proven success eliciting antibody responses with high titers and nanomolar binding affinities to carfentanil, fentanyl, and related synthetic opioids (Figure 1).^18^ The Carfen-*ester*-KLH immunoconjugate was formulated with our optimal adjuvant system consisting of CpG ODN 1826 and alum and vaccine was administered intramuscular (IM) on weeks 0, 4, 8 and 12 (Figure 2A). The immune response was monitored by Surface Plasmon Resonance (SPR) throughout the vaccination schedule and binding affinities for both carfentanil and fentanyl were measured for serum antibodies from week 6, 10, and 14 bleeds. A final IV boost consisting of immunoconjugate alone was administered 3 days prior to spleen and lymph nodes being harvested at week 16 (n=2 rats) for processing and B cell sorting. ^17^

Drug-specific B cell sorting was performed with a combination of carfentanil-and fentanyl-based probes with the intent to not only capture carfentanil-specific mAbs but also increase the pool to include fentanyl binders as well as pan-specific mAbs with broad cross-reactivity (Figure 1, Supplemental Figure S1). Complexing each probe with a distinct streptavidin-fluorophore conjugate through a biotin moiety strategically placed on the drug molecule provided a set of drug-specific fluorescent probes that were utilized for cell staining and B cell sorting. At the end of the sorting process, IgG^+^ single B cells positive for carfentanil and fentanyl were collected into individual wells of 96 well plates for a total of 288 single B cells. Next, single B cell RT-PCR followed by cloning into human mAb expression vectors (IgG1 heavy chain (HC) and appropriate kappa (κ) or lambda (λ) light chain (LC)) afforded 146 potential mAb clones with complete and in frame antibody genes consisting of cognate HC and LC pairs. The corresponding fully human mAbs were then expressed by transiently transfected HEK293 cells and culture supernatants containing individual mAbs were harvested and used directly for initial binding studies, initiating the mAb selection phase of our program.

SPR-based screening was used to assess the pool of prospective anti-opioid mAbs with respect to binding as well as cross-reactivity. All 146 candidates were first examined simply for their ability to bind the pertinent carfentanil-and/or fentanyl-BSA conjugates immobilized on the sensor chip surface, resulting in a binding test positivity rate of approximately 80%.^18-19^ This subset of binders (118) was next tested against a panel of synthetic opioids (carfentanil, fentanyl, acetylfentanyl, butyrylfentanyl, p-tolyfentanyl, 3-methylfentanyl, α-methylfentanyl, alfentanil and remifentanil) using SPR competitive assays. The resulting binding data were then scrutinized for drug affinity, specificity and broad cross-reactivity from which IC_50_ values were estimated and mAbs ranked (Supplemental Table S1). Based on a collective assessment of our initial findings, a set of top candidates showing high affinity for carfentanil and fentanyl along with pan-specificity were selected for further detailed characterization.

Before undergoing subsequent rounds of testing, lead mAbs were expressed and purified on small-scale. Kinetic analysis was then conducted using a Biacore 8K system with the purified mAbs immobilized on the chip surface and drug analyte passed over the antibody bound surface. Kinetic rate constants, association rate constant (*k*_*a*_) and dissociation rate constant (*k*_*d*_), were measured for both carfentanil and fentanyl and corresponding equilibrium dissociation constants (*K*_*D*_) calculated (Figure 2). Generally, all clones displayed subnanomolar to picomolar affinity forcarfentanil with exceptional *k*_*a*_ and *k*_*d*_ values along with a nanomolar to subnanomolar *K*_*D*_ range for fentanyl and desirable *k*_*a*_ and *k*_*d*_ values.

Encouraged by the binding results, we next used differential scanning calorimetry (DSC) and differential scanning fluorimetry (DSF) to measure the transition temperatures of the mAbs. These two complementary biophysical approaches enabled us to probe protein stability and aggregation propensity which can directly impact *in vivo* efficacy and t_1/2_. The thermal transition midpoints (*T*_m_) for each mAb were measured and all mAbs had an onset of melting temperature ≥ 62 °C in the absence of bound drug with values consistent across methods (Supplemental Table S2). Although the *T*_m_’s may be considered modest, the presence of carfentanil, i.e. mAb:carfentanil complex, substantially increased the *T*_m_ with the greatest shift observed for P1A4 in DSF experiments, a 15°C increase from 66°C to 81°C.

Considering the results of our binding studies, DSC and DSF analysis, and preliminary *in vivo* testing, the lead mAb selected to move forward for further development at this juncture was P1A4. This mAb displays excellent binding kinetics with a fast *k*_*a*_,slow *k*_*d*_,high affinity to carfentanil and fentanyl, and broad cross-reactivity with other synthetic opioids; however, improvement in its thermostability was warranted.

Phage display is a powerful and versatile technique that can be tailored for selection of desired properties and has proven invaluable in the development of therapeutic mAbs.^20^Taking advantage of the utility of phage display, we constructed mutant antibodies libraries using P1A4 in a single-chain variable fragment (scFv) format consisting of the variable heavy chain (VH) and variable light chain (VL) domains joined with a flexible (GGGS)_3_ linker. ScFv libraries were constructed using two different approaches and the resulting libraries were incorporated into our pIX phage display system for selection.^21^The first approach involved redesigning the antibody surface to increase the net charge of the protein in a process referred to as supercharging.^22^The second approach involved targeted random mutagenesis of selected framework residues. Using the initial crystal structure of wild-type P1A4, a scFv protein model was created and used as a guide for library design. Two separate scFv-phage libraries, reengineered surface library incorporating mutations at predicted surface residues and framework mutant library, were constructed and subjected to multiple rounds of panning against our carfentanil hapten with increasing stringency and elevated temperatures. Selected phage clones were analyzed by SPR and the most promising clones from this group underwent secondary analysis to assess binding as well as thermal stability.

Based on these results, residues that are predicted to have the most favorable impact on stability while maintaining binding kinetics and affinity were identified, one residue in the VL and one in the VH. Double mutant focused phage libraries were then generated targeting the VL residue in combination with the VH residue using NNY codon randomization for these positions. Library panning, selection, and analyses were performed *vide supra* with one clone clearly emerging as the best, containing an Ile to Asn mutation in the VH and Met to Ser in the VL. A third mutation was introduced to remove a potential glycosylation site within the VL at position Ser66 that was revealed in the initial crystal structure of wild-type P1A4 bound with carfentanil. Minimizing glycosylation within therapeutic mAbs is generally desired, thus this residue was mutated to a lysine residue to reduce the potential sequence liability moving forward. The purified scFv mutant displays much greater thermal stability (23°C increase in T_m_) and less aggregation propensity (5°C increase in T_agg_) compared to wild-type P1A4, while maintaining binding parameters. Our optimized mAb termed C10-S66K containing a total of 3 mutations (1 in the VH and 2 in the VL) was considered superior, therefore advancing in our program. The C10-S66K mAb displays high binding affinity (*K*_*D*_) for carfentanil, fentanyl with cross-reactivity to a panel of related synthetic opioids (Table 1). Moreover, there is little affinity to the clinically approved opioid analgesics alfentanil and remifentanil implying these two drugs could be used concomitantly with our antibody. Also noteworthy is a lack of affinity against the FDA-approved treatment for opioid overdose, naloxone.

**Table 1.** C10-S66K Binding Affinities

| Synthetic Opioid | $K_D$ (M) |
| --- | --- |
| Carfentanil | 6.899E-11 |
| Fentanyl | 2.277E-10 |
| Acetylfentanyl | 1.903E-10 |
| Butyrylfentanyl | 1.323E-10 |
| p-Tolylfentanyl | 1.781E-10 |
| 3-Methylfentanyl | 8.809E-09 |
| α-Methylfentanyl | 7.102E-10 |
| Sufentanil | 1.909E-08 |
| Alfentanil | >10 μM |
| Remifentanil | >1 μM |
| Naloxone | No binding |

The crystal structures of C10-S66K:fentanyl, C10-S66K:carfentanil, and apo C10-S66K were determined at resolutions of 1.80 Å, 1.78 Å, and 2.83 Å, respectively (Supplemental Table S3). Analysis of the complexes revealed that C10-S66K binds to fentanyl and carfentanil primarily through interactions with the phenylethyl and piperidinyl groups found in both compounds. These groups are deeply embedded in the binding pocket and form the core of the binding interaction (Figure 3A, Supplemental Figure S2A). The binding is predominantly driven by van der Waals forces, although a single hydrogen bond is formed between E50^H^ and the nitrogen in the piperidinyl ring of each compound (Figure 3B-C). Notably, aromatic residues F95^L^ and W34^H^ create a tight pocket that encloses the piperidinyl ring and locks in the drug. Since fentanyl has a pKa of 8.12 at human body temperature,^23^ its two nitrogens are protonated, driving favorable electrostatic interactions with the negatively charged pocket in C10-S66K (Supplemental Figure S3A). Interestingly, the conformations of the antibody complementarity-determining regions (CDRs) in both structures are identical (Supplemental Figure S3B), and in fact, C10-S66K employs the same residues to bind to both fentanyl and carfentanil (Figure 3B-C). Despite the overall similarities, carfentanil exhibits a slightly different conformation of its ketone group (Figure 3B, black arrow), which interacts with Y52^H^. However, the methoxycarbonyl group of carfentanil does not contribute any additional interactions.

**Figure 3.**
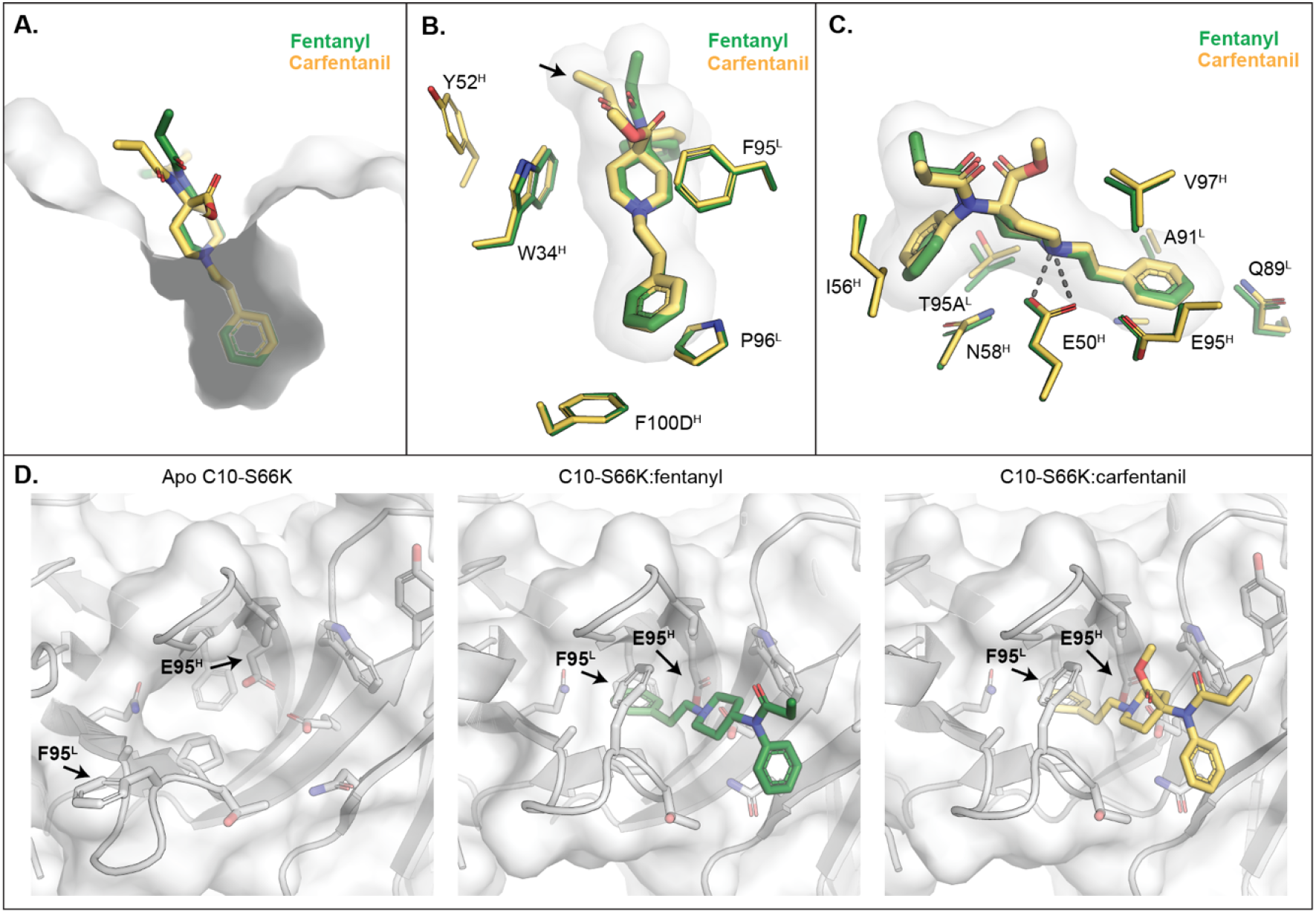
Crystal structures of C10-S66K. (A) Binding pocket of C10-S66K with fentanyl and carfentanil is displayed as a grey surface with fentanyl and carfentanil shown as green and yellow sticks, respectively. (B) Interactions of the drugs with aromatic residues from C10-S66K. Fentanyl and side chains from the C10-S66K:fentanyl complex are shown as green sticks, whereas those of carfentanil are in yellow. Fentanyl and carfentanil surfaces are also shown in light grey. (C) Interactions of the compounds with other residues in C10-S66K are displayed as in (B). (D) Comparison of residues in the paratope of C10-S66K with and without the drugs. Surfaces of C10-S66K of the corresponding complex are shown in grey. Stick representations of the fentanyl and carfentanil are shown in green and yellow, respectively. Black arrows indicate residues that display conformational changes.

Comparison of the apo C10-S66K Fab structure with the drug-bound complexes reveals conformational differences in two residues (Figure 3D). The side chain of E95^H^ moves inward to accommodate the phenylethyl ring of each compound, while the pocket for the drug appears wider in the apo structure due to a different conformation of F95^L^ (Figure 3D). Additionally, the side chain of F95^L^ rotates to clamp onto the piperidinyl ring of fentanyl/carfentanil (Figure 3D). It should be noted that the electron density of the side chain of F95^L^ in the apo structure is not fully resolved at a σ = 1 level, and the placement of the side chain is based on the electron density at a lower σ = 0.4 contour level (Supplemental Figure S3C). Thus, the electron density suggests that F95^L^ is flexible and that the pocket adopts a more open conformation in the absence of ligand and closes upon ligand binding.

Since C10-S66K predominantly interacts with the core phenylethyl and piperidinyl group in fentanyl/carfentanil, the antibody can bind to various fentanyl analogues sharing a similar backbone structure (Table 1, Supplemental Figure S2). The drugs align well based on the core groups, with differing conformations only in functional groups branching off the center nitrogen. The pocket of C10-S66K appears to be flexible enough to accommodate the ethylcyclopentane ring in sufentanil and the extra methyl group from the phenylethyl ring in α-methylfentanyl (Supplemental Figure S2). It is important to highlight that the additional functional groups branching from the piperidinyl ring in alfentanil and remifentanil may not fit as well into the pocket compared to carfentanil, as their orientations point in the opposite direction, facing the wall of the pocket. These structural differences, along with the substitution of the phenylethyl ring with a different structure, could contribute to the lower binding affinity observed for alfentanil and remifentanil (Table 1). Additionally, some high-affinity drugs have functional groups branching from the center N that may clash with CDR H2 in the bound C10-S66K structure, but the binding observed in Table 1 suggests that CDR H2 should be able to adopt different conformations to accommodate them (Supplemental Figure S2). In contrast, other published antibodies like HY6-F9^24^, FenAb609^12^, and FenAb208^12^have binding pockets that interact with unique functional groups of fentanyl, making them unlikely to bind to fentanyl analogs. Therefore, C10-S66K represents a promising candidate for antibody therapeutics that recognize fentanyl, carfentanil, and related drugs that share the core phenylethyl and piperidinyl group.

Next, the C10-S66K mAb in its minimal scFv format was evaluated *in vivo* to provide insight into the mAb’s therapeutic potential. To conduct these studies, the gene encoding the C10-S66K scFv was cloned into a mammalian expression vector and used to transiently transfect ExpiCHO suspension cells. Initially, a small batch of expressed protein was purified and administered to mice to determine its pharmacokinetic (PK) parameters. Mice received a 5 mg/kg dose of C10-S66K scFv (IP) and blood was collected over the course of 24 h. The amount of active antibody in plasma was determined by SPR and mAb concentrations were plotted against time to generate PK curves (Figure 4A). As evident, the scFv has a relatively short half-life (t_1/2_) of ∼0.9 h with an area under the curve (AUC) of 13 µg mL_-1_ h_-1_. The rapid clearance of smaller antibody fragments such as the scFv can be an advantage in some instances of acute drug overdose promoting swift renal excretion of the drug.^10, 25^ However, in the case of carfentanil, its pharmacological complexities (nonlinear accumulation and slow release) along with the potential for renarcotization following treatment with short-acting agents like naloxone led us to explore alternative means to better align the pharmacokinetic and pharmacodynamics of our mAb with carfentanil.^26^

**Figure 4.**
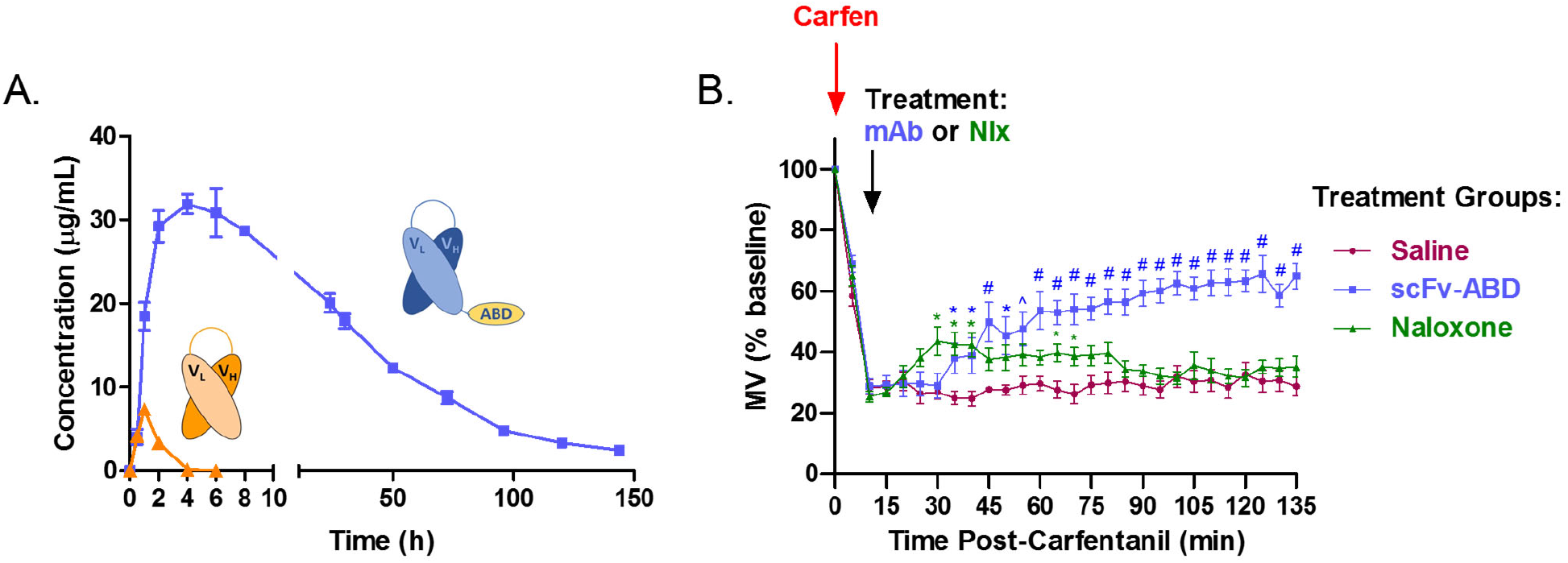
*In vivo* evaluation of C10-S66K monoclonal antibody. (A) Pharmacokinetic properties of scFv and scFv-ABD antibodies in mice injected intraperitoneally (IP) with 5mg/kg antibody. Blood samples were collected up to 6 days and active plasma antibody concentrations were determined surface plasmon resonance. (B) Reversal of carfentanil respiratory depression by C10-S66K scFv-ABD and naloxone. Mice (n = 8 per group) received a respiratory depressant dose of carfentanil (30 mg/kg, IP) at t=0 followed by treatment IP at t=15 min (30 mg/kg antibody, 1 mg/kg naloxone or saline). Respiration was measured by whole-body plethysmography and respiratory effects are plotted as percent of baseline minute volume (MV), established for 20 min prior to opioid administration. Data points denote the means ± SEM. Statistical comparison was made by two-way RM ANOVA to confirm a significant effect of treatment conditions [F (2, 21) = 14.81; p < 0.0001] with Bonferroni’s comparison; *p > 0.05, ^p > 0.01, #p > 0.001.

We previously reported the use of an albumin binding domain (ABD) as a means to increase the t_1/2_ of our nicotine degrading enzyme NicA2.^27^ Appendage of a 50 amino acid moiety (ABD035) to NicA2, extended the *in vivo* t_1/2_ of the fusion protein from ∼1 to 5 days. Therefore, a similar approach was explored for increasing the t_1/2_ of C10-S66K scFv. As illustrated in Figure 4A, addition of the ABD moiety markedly prolonged the *in vivo* t_1/2_ of the scFv from 0.9 to 47 h. Additionally, absorption of the scFv-ABD fusion mAb into the blood was greatly enhanced compared to scFv molecule resulting in a greater than 120-fold increase in AUC (1582 µg mL^-1^ h^-1^).

The improved scFv, C10-S66K scFv-ABD, with a suitable *in vivo* t_1/2_ was then tested in our respiratory depression mouse model using whole body plethysmography. Carfentanil displays dose-dependent respiratory depressive effects causing a measurable decrease in respiratory minute volume (MV), a product of tidal volume and respiratory rate.^18, 28^ Typically, a 30 µg/kg dose (IP) produces a ∼70% reduction in MV at maximum respiratory depression, 15 min post drug administration. In prior studies, we have demonstrated blockade of carfentanil’s respiratory depressive effects by vaccine elicited serum antibodies as well as passive immunization strategies, however a more pertinent model for counteracting drug intoxication as a countermeasure would be to demonstrate reversal of drug-induced respiratory depression post-exposure.^18, 28^ Therefore, in our current study mice were first dosed with 30 µg/kg carfentanil at t=0 followed by treatment at t=15 min (Figure 4B). Carfentanil dosed mice receiving C10-S66K scFv-ABD, 30 mg/kg, showed significant signs of recovery apparent at ∼20 min after mAb administration compared to the control group receiving saline. For comparison, the opioid antagonist naloxone which is currently the first line of defense against an opioid overdose was tested against an acute dose of carfentanil. This treatment group received a 1 mg/kg dose of naloxone which resulted in rapid reversal of carfentanil-induced respiratory depression although the beneficial effect is modest at best with signs of renarconization. In contrast, the mAb treatment group showed no indication of renarconization with sustained recovery.

The misuse of highly potent synthetic opioids has fueled the latest wave of the opioid crisis. Their illicit use for pain management or recreational purposes, along with potential weaponization continues to be an ongoing public health threat with the potential for global repercussions. The primary defense against opioid-related overdose is the short acting pharmaceutical naloxone that has proven less effective in treating the effects of ultrapotent synthetic opioids. Use of a sequestering agent such as a mAb provides an alternative and possible complementary approach for reversing opioid-induced overdose with the advantage of prolonged therapeutic efficacy.

With this purpose in mind, we isolated a pan-specific human mAb through drug-specific B cell sorting, then further improved its thermostability through phage display using a series of scFv phage libraries. This optimized mAb displays tight binding to a panel of fentanyl analogs with highest affinity for carfentanil, though binding affinities for other analogs are generally in the nanomolar range. In its minimal format, our mAb was able to reverse carfentanil-induced respiratory effects validating its utility against one of the most ultrapotent synthetic opioids. Importantly, no cross-reactivity was observed for naloxone supporting the concept of a potential combination therapy using a mAb and naloxone. While the drug landscape continues to change with many considering the new polysubstance trend a fourth wave, in most instances a synthetic opioid is involved. In these scenarios, for example xylazine contaminated with fentanyl, the recommended course of treatment is naloxone which treats the opioid-related pharmacology highlighting the ongoing need for effective remedies to counteract opioid-related overdose.

## METHODS

Details of experimental procedures are provided in the Supporting Information.

## ASSOCIATED CONTENT

### Supporting Information

Experimental procedures, probe synthesis, biochemical and *in vivo* procedures, X-ray crystallography and structural analysis, and supplemental tables and figures (PDF).

## Author Contributions

The manuscript was written through contributions of all authors. All authors have given approval to the final version of the manuscript.

## Notes

We thank Devan Sok and Elise Landais of the IAVI Neutralizing Antibody Center at Scripps Research for B cell sorting and monoclonal antibody cloning. We also thank Amanda Roberts of the Scripps Animal Behavioral Core for Immunizations and Tissue Collection. The antibodies in this manuscript are owned by TSRI and have been licensed to Cessation Therapeutics.

## ACKNOWLEDGMENT

Research reported in this manuscript was supported by the National Institute on Drug Abuse under Award No. U01DA046232.

