## Supporting Information for "An Engineered Human-Antibody Fragment with Fentanyl Pan-Specificity that Reverses Carfentanil-Induced Respiratory Depression"

### Table of Contents for Supporting Information


| | | ***Page*** |
|---|---|---|
| I. | Experimental Procedures and Biotin Probes | S2-S4 |
| | **Figure S1.** Carfentanil and Fentanyl Biotin Probes | S2 |
| II. | Immunization and Monoclonal Antibody Generation | S4 |
| III. | Biochemical Procedures | S4-S6 |
| IV. | X-ray Crystallography and Structural Analysis | S6 |
| V. | *In vivo* Studies | S7 |
| VI. | Supplemental Figures and Tables | S8-S13 |
| | **Table S1.** Binding Data and Cross-Reactivity of Monoclonal Antibodies | S8 |
| | **Table S2.** DSC and DSF Measurements of Selected Monoclonal Antibodies | S9 |
| | **Table S3.** X-ray data collection and refinement statistics | S10 |
| | **Figure S2.** Structural Analysis of C10-S66K and Predicted Binding Interactions | S11 |
| | **Figure S3.** Electrostatic Surface, CDRs, and Electron Density from C10-S66K Paratope | S12 |
| | **Table S4.** Biophysical Profile of scFv antibodies | S13 |
| | **Table S5**. Binding parameters of C10-S66K monoclonal antibody | S13 |
| | **Table S6**. Pharmacokinetic Parameters of C10-S66K scFv antibodies | S13 |
| VII. | References | S14 |

### I. Experimental Procedure

**Materials and Methods.** All reagents were purchased from commercial suppliers and used without further purification, unless otherwise stated. All reactions were carried out under argon unless otherwise noted. Anhydrous dichloromethane was distilled from calcium hydride under positive pressure of nitrogen. Preparative TLC separations were performed on Merck analytical plates (0.50 mm thick) precoated with silica gel 60 F254. Flash column chromatography was performed using the TELEDYNE ISCO (Combiflash Rf+) Lumen column chromatography system with silica gel grade, 230-400 mesh, 60 Å. Nuclear magnetic resonance $^{1}H$ and $^{13}C$ spectra were recorded on a Bruker DRX-600 or a Bruker DRX-500. Carfentanil and fentanyl probes (**1-4**) showed ≥95% purity by high-performance liquid chromatography (HPLC).

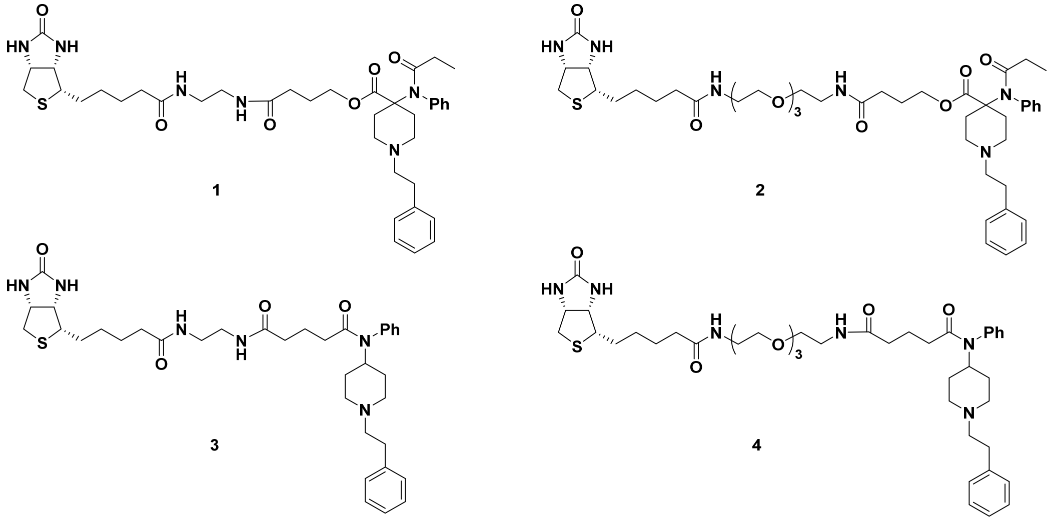

**Figure S1.** Carfentanil and fentanyl biotin probes.

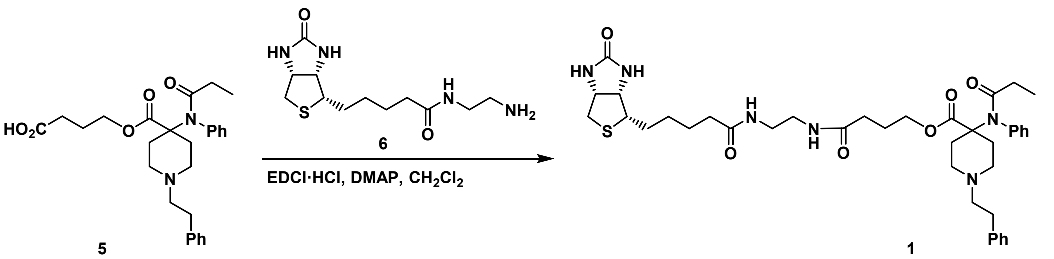

*4-oxo-4-((2-(5-((3aS,4S,6aR)-2-oxohexahydro-1H-thieno[3,4-d]imidazol-4-yl)pentanamido)ethyl)amino)butyl 1-phenethyl-4-(N-phenylpropionamido)piperidine-4-carboxylate (**1**).* Carfentanil derivative **5** and biotin derivative **6** were prepared according to known procedures.[1-2] To a solution of **5** (23.8 mg, 0.0511 mmol), DMAP (6.2 mg, 0.0511 mmol) and **6** (16.1 mg, 0.0562 mmol) in $CH_2Cl_2$ (0.8 mL) was added EDCI·HCl (13.7 mg, 0.0715 mmol) at 0 °C. The reaction mixture was stirred at 0 °C for 30 min followed by room temperature for 6 h. The mixture was diluted with $CH_2Cl_2$ (10 mL) and the solution was transferred to separating funnel. The organic layer was washed with saturated $NaHCO_3$ aq. (5 mL) and dried with $Na_2SO_4$. After filtration and evaporation, the crude was purified by silica gel chromatography ($CH_2Cl_2$/MeOH = 19/1 to 9/1) to give compound **1** as a pale-yellow oil (30.1 mg, 80%). $^{1}H$ NMR (600 MHz, Chloroform-*d*) δ 7.49 – 7.40 (m, 4H), 7.33 – 7.28 (m, 2H), 7.28 – 7.22 (m, 3H), 7.21 – 7.12 (m, 3H), 7.02 (t, *J* = 5.1 Hz, 1H), 6.17 (s, 1H), 5.42 (s, 1H), 4.49 (dd, *J* = 7.8, 4.9 Hz, 1H), 4.34 – 4.28 (m, 1H), 4.25 (t, *J* = 5.7 Hz, 2H), 3.45 – 3.36 (m, 4H), 3.18 – 3.11 (m, 1H), 2.92 – 2.85 (m, 1H), 2.80 – 2.71 (m, 4H), 2.63 – 2.40 (m, 4H), 2.33 – 2.24 (m, 6H), 2.13 – 2.05 (m, 2H), 1.93 (q, *J* = 7.4 Hz, 2H), 1.80 – 1.61 (m, 6H), 1.52 – 1.41 (m, 2H), 0.95 (t, *J* = 7.4 Hz, 3H). $^{13}C$ NMR (151 MHz, Chloroform-*d*) δ 175.18, 173.76, 173.73, 173.51, 163.74, 138.87, 130.42, 129.81, 129.28, 128.76, 128.56, 126.28, 63.37, 63.08, 61.83, 60.31, 55.60, 49.93, 40.67, 40.06, 39.81, 35.93, 33.55, 32.50, 29.84, 29.40, 28.23, 28.17, 25.60, 25.48, 9.33. HRMS-ESI: calculated for $C_{39}H_{55}N_6O_6S$ (M + H$^+$) 735.3898; found 735.3910.

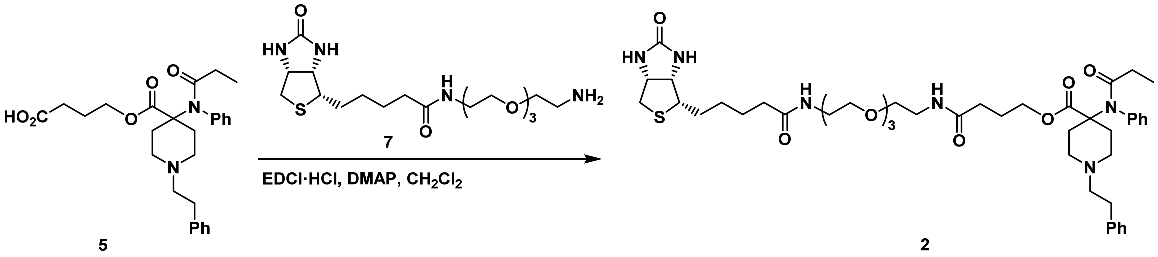

*4,18-dioxo-22-((3aS,4S,6aR)-2-oxohexahydro-1H-thieno[3,4-d]imidazol-4-yl)-8,11,14-trioxa-5,17-diazadocosyl 1-phenethyl-4-(N-phenylpropionamido)piperidine-4-carboxylate (**2**)*. Biotin derivative **7** was prepared according to known procedure.[3] To a solution of **5** (6.8 mg, 0.0146 mmol), DMAP (1.8 mg, 0.0146 mmol) and **7** (7.3 mg, 0.0175 mmol) in $CH_2Cl_2$ (0.5 mL) was added EDCI·HCl (3.9 mg, 0.0204 mmol) at 0 °C. The reaction mixture was stirred at 0 °C for 30 min followed by room temperature for 12 h. The mixture was diluted with $CH_2Cl_2$ (10 mL) and the solution was transferred to separating funnel. The organic layer was washed with saturated $NaHCO_3$ aq. (3 mL) and dried with $Na_2SO_4$. After filtration and evaporation, the crude was purified by preparative TLC ($CH_2Cl_2$/MeOH = 9/1) to give compound **2** as a pale-yellow oil (5.7 mg, 32%). $^1H$ NMR (600 MHz, Chloroform-*d*) δ 7.52 – 7.46 (m, 2H), 7.46 – 7.40 (m, 1H), 7.36 (d, *J* = 8.0 Hz, 2H), 7.33 – 7.27 (m, 2H), 7.26 – 7.18 (m, 3H), 7.17 – 7.04 (m, 1H), 4.55 – 4.18 (m, 3H), 3.68 – 3.56 (m, 10H), 3.56 – 3.53 (m, 2H), 3.51 – 3.34 (m, 6H), 3.30 – 3.09 (m, 7H), 2.89 (dd, *J* = 13.0, 4.9 Hz, 1H), 2.75 (d, *J* = 12.8 Hz, 1H), 2.52 – 2.33 (m, 8H), 2.33 – 2.17 (m, 4H), 2.10 – 2.02 (m, 2H), 1.93 (q, *J* = 7.4 Hz, 2H), 1.81 – 1.61 (m, 4H), 1.50 – 1.39 (m, 2H), 0.96 (t, *J* = 7.4 Hz, 3H).$^{13}C$ NMR (151 MHz, Chloroform-*d*) δ 175.18, 173.56, 172.63, 172.54, 138.06, 136.13, 130.33, 130.14, 129.67, 129.10, 128.84, 127.41, 70.52, 70.23, 70.19, 70.12, 70.05, 69.96, 64.90, 62.01, 60.52, 60.44, 58.26, 55.62, 49.81, 40.62, 39.36, 39.28, 35.81, 32.63, 30.52, 30.38, 29.11, 28.15, 28.13, 25.66, 25.07, 9.36. HRMS-ESI: calculated for $C_{45}H_{67}N_6O_9S$ (M + $H^+$) 867.4685; found 867.4685.

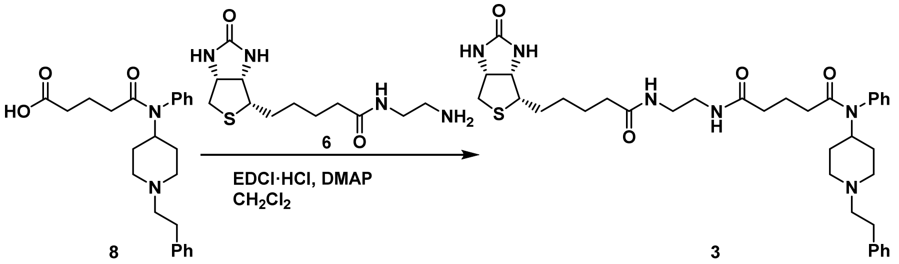

*N[1]-(2-(5-((3aS,4S,6aR)-2-oxohexahydro-1H-thieno[3,4-d]imidazol-4-yl)pentanamido)ethyl)-N[5]-(1-phenethylpiperidin-4-yl)-N[5]-phenylglutaramide (**3**)*. Fentanyl derivative **8** was prepared according to known procedure.[4] To a solution of **8** (16.5 mg, 0.0418 mmol), DMAP (5.1 mg, 0.0418 mmol) and **6** (13.2 mg, 0.0460 mmol) in $CH_2Cl_2$ (0.8 mL) was added EDCI·HCl (11.2 mg, 0.0586 mmol). The reaction mixture was stirred at room temperature for 10 h. After evaporation, the crude was purified by preparative TLC ($CH_2Cl_2$/MeOH 6/1) to give compound **3** as a colorless oil (11.8 mg, 43%).$^1H$ NMR (600 MHz, Methanol-$d_4$) δ 7.53 – 7.42 (m, 3H), 7.28 – 7.19 (m, 4H), 7.18 – 7.12 (m, 3H), 4.64 – 4.54 (m, 4H), 4.49 (dd, *J* = 7.9, 4.2 Hz, 1H), 4.31 (dd, *J* = 7.9, 4.5 Hz, 1H), 3.24 – 3.18 (m, 5H), 3.09 – 3.02 (m, 2H), 2.92 (dd, *J* = 12.8, 5.0 Hz, 1H), 2.76 – 2.68 (m, 3H), 2.57 – 2.50 (m, 2H), 2.25 – 2.14 (m, 4H), 2.09 (t, *J* = 7.4 Hz, 2H), 2.00 (t, *J* = 7.4 Hz, 2H), 1.88 – 1.55 (m, 9H), 1.50 – 1.39 (m, 4H). $^{13}C$ NMR (151 MHz, Methanol-$d_4$) δ 176.35, 175.74, 174.19, 166.09, 141.11, 139.69, 131.50, 130.69, 129.92, 129.62, 129.47, 127.17, 63.36, 61.62, 61.32, 56.97, 53.96, 53.85, 41.06, 40.02, 39.95, 36.82, 36.22, 35.26, 34.07, 31.12, 29.78, 29.49, 26.78, 22.70. HRMS-ESI: calculated for $C_{36}H_{51}N_6O_4S$ (M + $H^+$) 663.3687; found 663.3687.

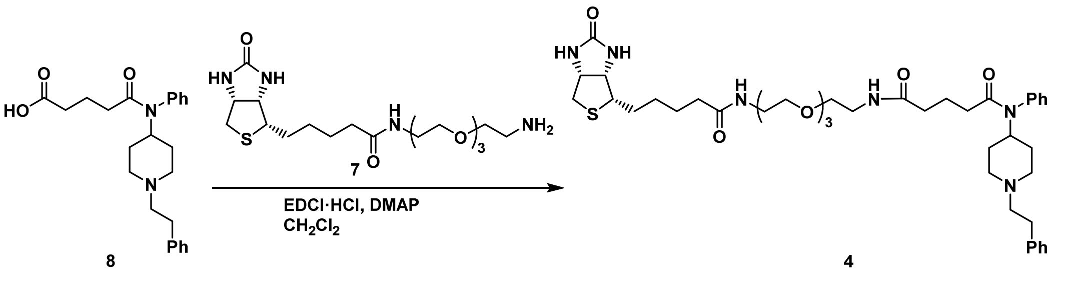

*N[1]-(13-oxo-17-((3aS,4S,6aR)-2-oxohexahydro-1H-thieno[3,4-d]imidazol-4-yl)-3,6,9-trioxa-12-azaheptadecyl)-N[5]-(1-phenethylpiperidin-4-yl)-N[5]-phenylglutaramide (**4**)*. To a solution of **8** (12.0 mg, 0.0304 mmol), DMAP (3.7 mg, 0.0304

mmol) and biotin derivative **7** (14.0 mg, 0.0334 mmol) in $CH_2Cl_2$ (0.6 mL) was added EDCI·HCl (8.2 mg, 0.0426 mmol). The reaction mixture was stirred at room temperature for 10 h. After evaporation, the crude was purified by preparative TLC ($CH_2Cl_2$/MeOH 6/1) to give compound **4** as a colorless oil (10.8 mg, 45%). $^1H$ NMR (600 MHz, Methanol-$d_4$) δ 7.53 – 7.43 (m, 3H), 7.28 – 7.19 (m, 4H), 7.19 – 7.13 (m, 3H), 4.64 – 4.56 (m, 5H), 4.52 – 4.46 (m, 1H), 4.30 (dd, $J$ = 7.9, 4.4 Hz, 1H), 3.67 – 3.61 (m, 6H), 3.61 – 3.57 (m, 2H), 3.54 (t, $J$ = 5.5 Hz, 2H), 3.48 (t, $J$ = 5.6 Hz, 2H), 3.37 – 3.35 (m, 2H), 3.29 (t, $J$ = 5.6 Hz, 2H), 3.24 – 3.17 (m, 1H), 3.09 (d, $J$ = 12.2 Hz, 2H), 2.92 (dd, $J$ = 12.7, 5.0 Hz, 1H), 2.78 – 2.68 (m, 3H), 2.61 – 2.55 (m, 2H), 2.31 – 2.20 (m, 4H), 2.11 (t, $J$ = 7.4 Hz, 2H), 2.00 (t, $J$ = 7.4 Hz, 2H), 1.92 – 1.55 (m, 8H), 1.51 – 1.40 (m, 4H). $^{13}C$ NMR (151 MHz, Methanol-$d_4$) δ 176.12, 175.42, 174.22, 166.09, 140.90, 139.69, 131.51, 130.72, 129.94, 129.63, 129.50, 127.24, 71.59, 71.57, 71.22, 70.60, 70.53, 63.37, 61.62, 61.18, 57.00, 53.94, 53.69, 41.06, 40.34, 40.27, 36.75, 36.09, 35.23, 33.93, 31.00, 29.77, 29.51, 26.85, 22.75. HRMS-ESI: calculated for $C_{42}H_{63}N_6O_7S$ (M + $H^+$) 795.4473; found 795.4490.

### II. Immunizations and Monoclonal Antibody Generation

All animal procedures were performed under IACUC-approved protocols and guidelines. The Carfen-*ester*-KLH conjugate was formulated with CpG ODN 1826 and alum (Alhydrogel®, Invivogen).[1] Human IgG-transgenic rats (Omni Rats, n = 2) were immunized with a series of IM tail base injections consisting of 100 μg Carfen-*ester*-KLH, 100μg CpG ODN, 0.75mg alum per dose at weeks 0, 4, 8, 12 with a final IV boost of Carfen-*ester*-KLH alone 3 days prior to terminal procedures. Blood samples were collected at weeks 6, 10, 14. Spleen and lymph nodes were harvested at week 16 for B cell sorting.[5]

Drug-specific single B cell sorting, single cell RT-PCR and cloning were performed by the IAVI Neutralizing Antibody Center at Scripps Research.[5] Cell suspensions were prepared from spleen and iliac lymph nodes and used for single B cell sorting into 96 well plates. In brief, after gating for lymphocytes (FSC x SSC) and singlets (FSC (H) x FSC (W)), live cells (FVS510-) followed by B cells were identified. From this population anti-rat IgG (IgG1-2a-2b-FITC) stained cells were further sorted using a mixture of carfentanil and fentanyl biotin probes (**1-4**), each linked to a distinct streptavidin fluorophore. Cells dual positive for carfentanil probes (**1,2**) and/or fentanyl probes (**3,4**) were collected. The plates containing single B cell were then processed through single cell RT-PCR, cloning into human IgG1 heavy chain (HC) and appropriate kappa (κ) or lambda (λ) light chain (LC) expression vectors, and sequence verification.[5]

### III. Biochemical Procedures

#### Surface Plasmon Resonance.

*1. SPR Screening of potential monoclonal antibody (mAb) clones.* A. Binding of the 146 mAb candidates was determined on a Biacore 3000 (GE Healthcare) using a CM5 sensor chip at 25 °C using HBS-EP+ buffer as running buffer. The ligands, carfentanil-BSA and fentanyl-BSA conjugates, were immobilized individually on the chip surface using standard NHS/EDC coupling chemistry with BSA alone as the reference flow cell.[1, 4] Cell culture supernatants containing individual mAbs were injected for 5 min at a flow rate of 30 μl/min and the binding stability signal 2.5 min after the injection was recorded in the sensogram. The chip surface was then regenerated with Gly-HCl (pH1.5) for 30 s before the next cycle of analysis. Antibody binding signal that is greater than 10 x SD of running buffer signal was considered positive binding, resulting in 118 positive clones. B. Competitive assays with the subset of positive binders was conducted against a panel of synthetic opioids, including carfentanil, fentanyl, acetylfentanyl, butyrylfentanyl, p-tolyfentanyl, 3-methylfentanyl, α-methylfentanyl, alfentanil and remifentanyl. Compounds at a final concentration of 0, 1, 10, 100, and 1000 nM were incubated with mAb supernatants at room temperature for a minimum of 30 min before the mixture was passed over the chip surfaces as described *vide supra*. The $IC_{50}$ values were estimated based on the binding response (RU) at various compound concentrations compared to no compound samples (0 nM) for each drug set.

2. *Binding kinetic analysis of top mAb candidates*. Binding kinetic analysis of lead mAbs was conducted on a Biacore 8K (GE Healthcare) instrument equipped with a series S sensor chip CM5 at 25 °C using PBS buffer as running buffer at a flow rate of 30 μl/min. In brief, individual antibodies were immobilized on the chip surface in separate channels at a response level of ~2500 RU and human serum albumin (HSA) was immobilized on the reference flow cell of each channel at an equivalent level, all using standard NHS/EDC coupling chemistry. To determine the binding kinetics, a predefined single-cycle kinetics (SCK) protocol (LMW single-cycle kinetics, Biacore 8K Control Software ver. 4.0.8.19879) was used as follows: 1) 3 startup cycles (each cycle consists of a 120 s injection of running buffer and a 300 s dissociation phase followed by regeneration with Gly-HCl (pH1.5) for 30 s) were run before SCK analysis. 2) For SCK analysis, fentanyl, carfentanil and fentanyl derivatives were prepared in running buffer at concentrations of 3.75, 7.5, 15, 30, and 60 nM. Each compound dilution (from low to high) was passed over the flow cells (HSA reference then active antibody flow cells in each channel) for 50 s consecutively followed by 9800 s of dissociation in running buffer. 3) The sensor chip surface was regenerated with an injection of Gly-HCl (pH 1.5) solution for 30 s before the next cycle of SCK analysis. 4) Data collection rate of 10 Hz was used for all SCK analyses. A blank running buffer injection was also performed before

each compound injection as a reference using the same conditions for SCK analysis. The data collected by the Biacore 8K control software in results files were then analyzed by Biacore Insight evaluation software (ver. 4.0.8.19879). Each SCK analysis data set was double referenced with signal from the reference HSA flow cell and blank running buffer cycles and was fit using a 1:1 binding model.

2. *Binding kinetics of C10-S66K IgG*. The binding kinetics for C10-S66K IgG against carfentanil, fentanyl and various derivatives were measured on a Biacore S200 (GE Healthcare) instrument equipped with a series S sensor chip CM5 at 25 °C using PBS running buffer at a flow rate of 50 μl/min. C10-S66K IgG was immobilized on the active flow cell at a response level of ~2000 RU and HSA on the reference flow cell at an equivalent level, using standard NHS/EDC coupling chemistry. A predefined SCK protocol (LMW kinetics/affinity single cycle, Biacore S200 Control Software ver. 1.1 build 28) was used as follows: 1) 3 startup cycles (each cycle consists of a 60 s injection of running buffer and a 300 s dissociation phase followed by regeneration with Gly-HCl (pH1.5) for 30 s) were run before SCK analysis. 2) For SCK analysis, fentanyl, carfentanil and related drugs were prepared in running buffer at concentrations of 2.5, 5, 10, 20, and 40 nM. Each compound dilution (from low to high) was injected for 60 s through both reference and active flow cells consecutively followed by 10800 s of dissociation in running buffer. 3) The chip surface was regenerated by injection of a Gly-HCl (pH 1.5) solution for 30 s before the next cycle of SCK analysis. 4) Data collection rate of 10 Hz was used for all SCK analyses. A blank running buffer injection done prior to each compound injection as a reference. All data collected by Biacore S200 control software in a result file were analyzed by Biacore S200 evaluation software (ver. 1.1 build 27). Each SCK analysis data set was double referenced with signal from reference flow cell and blank running buffer injection and was fitted using a 1:1 binding model.

**Biophysical Measurements.**

*1. Thermal transition midpoints ($T_m$) measurements using Differential scanning calorimetry (DSC) and differential scanning fluorimetry (DSF).* A protein in solution is in equilibrium between its native and denatured conformations with a higher $T_m$ value reflecting a more stable protein. To evaluate the thermal stability of our top mAb candidates, DSC and DSF were used to measure $T_m$ values and compare thermal stability among the mAbs. For these measurements, antibody alone or complexed with carfentanil and matched buffer samples (reference) were prepared in triplicates and applied to MicroCal VP-Capillary DSC (Malvern Instruments, Northampton, MA) for DSC and Prometheus NT.48 (NanoTemper Technologies) for DSF analysis. All samples were heated from 25 °C to 100 °C at a ramp rate of 0.5 °C/min. Thermograms for each mAb or mAb complex was analyzed and $T_m$ values for each sample were calculated using corresponding instrument analysis software.

*2. Biophysical profiles or P1A4 scFv and C10-S66K scFv using Uncle System.* To evaluate the thermal stability, aggregation propensity and polydispersity of P1A4 scFv and C10-S66K scFv, we used the all-in-one Uncle System (Unchained Labs) to apply dynamic light scattering (DLS), static light scattering (SLS) and intrinsic protein fluorescence scanning. For these measurements, 9 μl of sample was loaded into the Uni in triplicate followed by docking of the Uni into the instrument for measurement. All samples were heated from 15 °C to 95 °C at a ramp rate of 0.5 °C/min. DLS was measured before and after the thermal ramp. The full emission spectrum from 250–720 nm was measured at every measured temperature. The thermogram was analyzed using Uncle analysis software (ver. 5.03). Polydispersity and hydrodynamic diameter (in nm) were measured by DLS before thermal ramp and fitted by the analysis software. The $T_{agg}$ was measured by $SLS_{266nm}/SLS_{473nm}$ and calculated for all samples by the analysis software. The $T_m$ was measured by intrinsic protein fluorescence changes during the entire thermal ramping process and analyzed using Barycentric mean method (BCM) in the analysis software.

**Phage Display**.

*1. Construction and panning of scFv supercharging library.* It has been reported that supercharging some proteins may significantly enhance their solubility and aggregation resistance.[6] Therefore, we anticipated that the supercharging method may improve the thermal stability and solubility of P1A4. First, we identified a few potential mutation residues in the scFv variable region of heavy and light chains using a supercharging application in Rosetta-mode with a target net charge with default setting (https://rosie.rosettacommons.org/supercharge/). The philosophy of Rosetta-mode supercharge is to mutate residues positions that preserve and/or add favorable surface interactions.[7] Residues H63S, H66S, H88A and H89A of the HC and residues L14S, L41D, L67S and L80A of the LC were selected for mutation. All residues were mutated to either lysine or arginine via site-directed mutagenesis using codon "ARA". The mutant scFv genes were then cloned into the pCGMT9 phage display vector, creating a focused mutant library. The scFv-supercharging mutant library was amplified and panned against carfentanil-BSA as described.[8] After two rounds of panning, 96 clones were randomly picked and phage-scFvs were amplified. The supernatants containing phage-scFv mutants were used to select binders against carfentanil-BSA by phage ELISA.[8]

*2. Construction and panning of scFv framework single mutant library.* Computational modeling of changes in binding free energies (ΔΔG) imposed by mutation(s) allows prediction and perturbation of protein-protein interactions and structure stability. Therefore, we sought to identify potential mutation(s) in the framework region that could strengthen interactions within VH and VL interface. First, we identified a panel of VH-VL interface residues using the initial wild-type P1A4 structure by running a "InterfaceResidues" script in PyMol. Next, the selected interface residues were subjected to $\Delta\Delta G$ calculations using the point mutant scan application (pmut_scan_parallel) from the Rosetta Software Suite (https://www.rosettacommons.org/software). According to the results, seven residue positions with a $\Delta\Delta G$ decrease between -0.5 and -5.0 kcal/mol were selected for mutation. To construct a single site mutation scFv library, each selected residue was randomly mutated using codon NNY and the pool of single mutant scFv genes were cloned into pCGMT9 phage display vector. The scFv-single mutant library was amplified and panned against carfentanil-BSA as described.[8] After four rounds of panning, 96 clones were randomly picked and phage-scFvs were amplified. The supernatant containing phage-scFv mutant was used to select binders against carfentanil-BSA by Biacore binding analysis *vide supra*. Phage-scFv mutants displaying a stronger binding response compared to wild-type P1A4 phage-scFv were selected to undergo further mutagenesis through generation of double mutant libraries.

*2. Construction and panning of scFv framework double mutant library.* A combination of two mutation positions of the selected single mutants was randomly mutated using codon NNY, i.e., two mutations per scFv gene, and the pool of double mutant scFv genes was cloned into pCGMT9 phage display vector. The scFv-double mutant library was amplified and panned against carfentanil-BSA as described.[8] After four rounds of panning, 96 clones were randomly picked and phage-scFvs were amplified. Supernatants containing phage-scFv mutants were used to select binders against carfentanil-BSA by Biacore binding analysis *vide supra*. Phage-scFv mutants displaying a stronger binding response compared to wild-type were selected for soluble scFv expression, purification and biophysical characterization.

**Mammalian Cell Expression and Purification of C10-S66K scFv and C10-S66K scFv-ABD**.

The C10-S66K scFv and scFv-ABD genes were codon optimized for mammalian cell expression and synthesized by GeneScript Biotech Corp. as an expression cassette with the following elements: 5' Hind III site followed by Kozak sequence and hLSP (MAWTPLFLLTCCPGSNS human lambda light chain signal peptide), then the His x 6 affinity tag followed by the C10-S66K scFv/scFv-ABD gene, and finally a 3' Xho I site. The complete expression cassette was excised from the cloning vector (GeneScript) with restriction enzymes Hind III and Xho I and ligated into the mammalian expression vector pcDNA 5/FRT/TO (Invitrogen, Cat. No. V652020) using vector multiple cloning sites Hind III and Xho I. The pcDNA 5/FRT/TO vector contains a hybrid human cytomegalovirus (CMV)TetO2 promoter that can be used for high-level, tetracycline-regulated expression of the gene of interest in mammalian cells and is compatible with the ExpiCHO expression system (ThermoFisher Scientific, Cat. No. A29133) chosen to produce the C10-S66K scFv and C10-S66K scFv-ABD proteins. In brief, ExpiCHO-S cells were transiently transfected with 0.5 mg plasmid/ml cells, and cultures were induced with 0.1 mg/mL doxycycline hyclate (Sigma, Cat. No. D5207) on day 3 post-transfection and the cells were grown using the max titer protocol per manufacturer's instruction. Cultures were harvested 12-14 days post-transfection or whenever the cell viability was dropped below 75%. The culture was clarified by centrifugation and the supernatant was filtered through a 0.22-mm filter. The protein was purified from the filtered supernatant by IMAC using His60 Ni superflow resin (TakaRa Bio, Cat. No. 635660). The purified protein, greater than 95% homogeneity by SDS-PAGE, was dialyzed against PBS and concentrated for biophysical characterization and *in vivo* studies.

### IV. X-ray Crystallography and Structural Analysis

Fab from antibody C10-S66K was mixed with fentanyl or carfentanil in a 1:1 molar ratio. For Fabs used in the fentanyl/carfentanil complex, six substitutions and one deletion (from $^{112}$SSASTKG$^{118}$ to $^{112}$VSRRLP$^{117}$) were introduced into the elbow region of the Fab to facilitate crystallization as previously described.[9] The apo C10-S66K Fab was mixed with Streptococcal immunoglobulin G-binding protein G (domain III) in the Fab to protein G ratio of 1:1 to promote crystallization of the Fabs.[10] Crystal screening of the complexes was performed using our high-throughput, robotic CrystalMation system (Rigaku, Carlsbad, CA) at The Scripps Research Institute, which was based on the sitting drop vapor diffusion method with 35 μL reservoir solution and each drop consisting of 0.1 μL protein + 0.1 μL precipitant. C10-S66K Fab:fentanyl co-crystals were grown in 1.0 M Li-chloride, 10% PEG-6000, 0.1 M Bicine pH 9.0 and were cryoprotected in 10% ethylene glycol. C10-S66K Fab:carfentanil co-crystals grew in 1.0 M Li-chloride, 10% PEG-6000, 0.1 M HEPES pH 7.0 and were cryoprotected in 15% ethylene glycol. Apo C10-S66K Fab crystals grew 20% PEG 3350, 0.2 M $Na_3$-citrate, pH 8.2 and were cryoprotected in 10% ethylene glycol. X-ray diffraction data were collected at the Advanced Proton Source (APS) beamline 23IDD, or at the Stanford Synchrotron Radiation Lightsource beamline 12-1, and processed and scaled using the HKL-2000 package.[11] The structures were obtained using molecular replacement with Phaser.[12] Refinement of the structures was carried out using phenix.refine[13] and involved multiple iterations with Coot.[14] The Kabat system was used to number the amino acid residues in the Fabs, and MolProbity[15] was used to validate the structures. For structural analysis, buried surface areas (BSAs) were calculated with the program MS,[16] and hydrogen bonds were assessed with the program HBPLUS.[17] The crystal structures of apo C10-S66K Fab and complexes with fentanyl and carfentanil have been deposited in the Protein Data Bank.

### V. *In vivo* Studies.

Female Swiss Webster mice, 6 weeks old, were obtained from Taconic Farms (Germantown, NY) and were acclimated for at least 1 week after arriving. Mice were housed at 22 °C on a reversed 12 h light cycle (9 p.m. to 9 a.m.) with *ad libitum* access to food and water. All experiments were performed in the dark (active) phase. All animal procedures were performed under IACUC-approved protocols and guidelines.

#### Pharmacokinetics of C10-S66K scFv and C10-S66K scFv-ABD.

Mice (n = 6 per group) were injected intraperitoneally with C10-S66K scFv or C10-S66K scFv-ABD at a dose of 5 mg/kg. Blood samples were taken via retro-orbital bleed at each time point to generate n = 2 for each measured timepoint, alternating the animals over the timepoints. Collected blood samples were then centrifuged at 10,000 rpm for 10 min and the serum was collected. The concentration of active mAb was determined by SPR using the Biacore 3000 method as described *vide supra*. The RU for mAb binding was measured 2.5 min after the end of each injection. Quantification of mAb plasma concentrations was achieved by including a set of mAb standards during each corresponding Biacore analysis, generating a standard curve (plot of mAb concentration against time). The elimination rate constant ($k_e$) for each mAb was calculated by plotting the natural log against time during the elimination phase and determining the slope. The half-life ($t_{1/2}$) was then calculated using the equation Ln(2)/$k_e$. (Supplemental Table 6).

#### Respiratory Depression Studies.

Respiration was measured in conscious freely moving mice whole-body plethysmography chambers (EMKA Technologies, France), as previously described.[1, 18] One day prior to the experiment, mice were habituated to the chambers for 30 min. On the experimental day, baseline measurements were recorded for 20 min before receiving a 30 µg/kg dose of carfentanil intraperitoneally (t = 0) within 5 min of baseline and returned to the chamber. At maximum respiratory depression (t = 15 min), mice received treatment intraperitoneally and respiration was measured for an additional 2 h. Treatment groups (n = 8/group) consisted of a C10-S66K scFv-ABD (30mg/kg), naloxone (1 mg/kg), and control saline group. Using IOX software by EMKA, data were recorded and binned into 5-min intervals. Changes in minute volume (MV) were used to assess changes in respiration and the percent change in MV was calculated for each mouse as the percent of pre-drug baseline.

#### Statistical Analysis.

All data are displayed as mean ± SEM. Data were graphed and analyzed using GraphPad Prism, setting $p < 0.05$ as the critical value. Respiratory depression statistical comparison was made by two-way RM ANOVA with Bonferroni's comparison.

### VI. Supplemental Figures and Tables

| mAb | Carfen | Fen | AcetylFen | ButyrylFen | TolyFen | 3-MethylFen | α-MethylFen | AlFen | RemiFen |
|---|---|---|---|---|---|---|---|---|---|
| P1A12 | < 1nM | > 10nM | < 10nM | < 10nM | < 10nM | < 10nM | > 10nM | >> 1μM | >> 0.1μM |
| P1A2 | < 1nM | < 1nM | < 10nM | < 10nM | ~ 10nM | > 10 nM | < 10nM | >> 1μM | >> 0.1μM |
| P1A4 | < 1nM | < 1nM | < 10nM | < 10nM | < 10nM | < 10nM | < 10nM | > 1μM | > 0.1μM |
| P1A5 | ~ 1nM | >> 10nM | >> 10nM | >> 10nM | >> 10nM | >> 10nM | >> 10nM | >> 1μM | < 0.1μM |
| P1A6 | ~ 1nM | >> 10nM | >> 10nM | >> 10nM | >> 10nM | >> 10nM | >> 10nM | < 1μM | < 0.1μM |
| P1A7 | < 1nM | < 1nM | < 10nM | < 10nM | < 10nM | > 10nM | < 10nM | < 1μM | < 0.1μM |
| P1A9 | < 1nM | >> 10nM | > 10nM | ~ 10nM | ~ 10nM | > 10nM | >> 10nM | >> 1μM | >> 0.1μM |
| P1B11 | < 10nM | < 10nM | ~ 10nM | < 10nM | >> 10nM | >> 10nM | < 10nM | >> 1μM | < 0.1μM |
| P1B3 | ~ 1nM | > 10nM | >> 10nM | ~ 10nM | >> 10nM | >> 10nM | < 10nM | < 1μM | < 0.1μM |
| P1B4 | < 10nM | >> 10nM | >> 10nM | >> 10nM | >> 10nM | >> 10nM | >> 10nM | ~ 1μM | < 0.1μM |
| P1B5 | < 1nM | >> 10nM | >> 10nM | >> 10nM | >> 10nM | >> 10nM | >> 10nM | < 1μM | < 0.1μM |
| P1B7 | ~ 1nM | > 10nM | ~10nM | ~10nM | >> 10nM | ~10nM | >> 10nM | >> 1μM | >> 0.1μM |
| P1B8 | < 10nM | >> 10nM | >> 10nM | >> 10nM | >> 10nM | >> 10nM | >> 10nM | < 1μM | < 0.1μM |
| P1C1 | < 1nM | > 10nM | < 10nM | < 10nM | < 10nM | ~10nM | > 10nM | >> 1μM | >> 0.1μM |
| P1C10 | ~1nM | >> 10nM | >> 10nM | >> 10nM | >> 10nM | >> 10nM | >> 10nM | >> 10μM | >> 0.1μM |
| P1C12 | < 1nM | < 10nM | < 10nM | < 10nM | ~ 10nM | < 10nM | ~ 10nM | >> 1μM | >> 0.1μM |
| P1C3 | ~ 1nM | > 10nM | ~ 10nM | < 10nM | >> 10nM | >> 10nM | >> 10nM | >> 1μM | >> 0.1μM |
| P1C8 | < 1nM | < 10nM | < 10nM | < 10nM | < 10nM | < 10nM | ~ 10nM | >> 1μM | >> 0.1μM |
| P1C9 | < 1nM | > 10nM | >> 10nM | >> 10nM | >> 10nM | >> 10nM | < 10nM | ~ 1μM | < 0.1μM |
| P1D1 | < 1nM | < 10nM | < 10nM | < 10nM | < 10nM | < 10nM | < 10nM | > 1μM | >> 0.1μM |
| P1D11 | < 10nM | < 10nM | < 10nM | < 10nM | >> 10nM | >> 10nM | < 10nM | >> 1μM | >> 0.1μM |
| P1D3 | < 1nM | < 10nM | < 10nM | < 10nM | >> 10nM | >> 10nM | > 10nM | > 1μM | >> 0.1μM |
| P1D8 | < 1nM | < 1nm | < 10nM | < 10nM | < 10nM | < 10nM | < 10nM | >> 1μM | >> 0.1μM |
| P1E10 | < 1nM | < 10nM | < 10nM | < 10nM | ~ 10nM | < 10nM | > 10nM | >> 1μM | >> 0.1μM |
| P1E11 | ~ 1nM | >> 10nM | >> 10nM | >> 10nM | >> 10nM | >> 10nM | >> 10nM | < 1μM | < 0.1μM |
| P1E12 | < 1nM | < 10nM | < 10nM | < 10nM | >> 10nM | >> 10nM | > 10nM | > 1μM | >> 0.1μM |
| P1E2 | < 1nM | < 10nM | < 10nM | < 10nM | >> 10nM | >> 10nM | < 10nM | ~ 1μM | < 0.1μM |
| P1E4 | ~ 1nM | < 10nM | < 10nM | < 10nM | < 10nM | >> 10nM | < 10nM | > 1μM | >> 0.1μM |
| P1E7 | < 1nM | < 1nM | < 10nM | < 10nM | < 10nM | < 10nM | < 10nM | >> 1μM | >> 0.1μM |
| P1E7 | < 1nM | < 1nM | < 10nM | < 10nM | < 10nM | < 10nM | < 10nM | >> 1μM | >> 0.1μM |
| P1F11 | < 1nM | < 1nM | < 10nM | < 10nM | < 10nM | > 10nM | < 10nM | > 1μM | > 0.1μM |
| P1F2 | < 10nM | < 10nM | ~ 10nM | ~ 10nM | ~ 10nM | >> 10nM | > 10nM | >> 1μM | >> 0.1μM |
| P1F5 | ~ 1nM | >> 10nM | >> 10nM | >> 10nM | >> 10nM | >> 10nM | >> 10nM | ~ 1μM | < 0.1μM |
| P1F8 | < 1nM | ~ 1nM | < 10nM | < 10nM | < 10nM | < 10nM | < 10nM | >> 1μM | >> 0.1μM |
| P1G1 | < 1nM | < 10nM | < 10nM | < 10nM | >> 10nM | >> 10nM | > 10nM | >> 1μM | >> 0.1μM |
| P1G12 | ~ 1nM | < 10nM | < 10nM | ~ 10nM | >> 10nM | ~ 10nM | ~ 10nM | >> 1μM | >> 0.1μM |
| P1H2 | < 1nM | < 10nM | < 10nM | < 10nM | > 10nM | >> 10nM | ~ 10nM | >> 1μM | > 0.1μM |
| P2A11 | < 1nM | < 10nM | < 10nM | < 10nM | > 10nM | >> 10nM | > 10nM | >> 1μM | >> 0.1μM |
| P2A7 | < 1nM | < 1nM | < 10nM | < 10nM | < 10nM | ~ 10nM | < 10nM | >> 1μM | >> 0.1μM |
| P2B5 | < 1nM | < 1nM | < 10nM | < 10nM | < 10nM | ~ 10nM | < 10nM | > 1μM | < 0.1μM |
| P2C1 | < 1nM | ~ 1nM | < 10nM | < 10nM | < 10nM | < 10nM | < 10nM | >> 1μM | >> 0.1μM |
| P2C12 | < 10nM | >> 10nM | >> 10nM | >> 10nM | >> 10nM | >> 10nM | >> 10nM | >> 1μM | < 0.1μM |
| P2C5 | ~ 1nM | < 10nM | < 10nM | < 10nM | > 10nM | >> 10nM | ~ 10nM | >> 1μM | >> 0.1μM |
| P2D10 | ~ 1nM | < 1nM | < 10nM | < 10nM | < 10nM | > 10nM | < 10nM | >> 1μM | ~ 0.1μM |
| P2D12 | < 10nM | < 10nM | < 10nM | < 10nM | < 10nM | >> 10nM | < 10nM | >> 1μM | >> 0.1μM |

>> no inhibition at all at tested compound concentration; > inhibits less than 50% of the binding at tested compound concentration; < inhibits more than 50% of the binding at tested compound concentration; ~ inhibits about 50% of the binding at tested compound

**Table S1.** Estimated $IC_{50}$ values for carfentanil, fentanyl, and other synthetic opioids.

| mAb | DSC | | | | DSF | | | DSF* | |
|---|---|---|---|---|---|---|---|---|---|
| | Tm1 | Tm2 | Tm3 | Tm4 | Tm1 | Tm2 | Tm3 | Tm1 | Tm2 |
| P1A4 | *66.70* | 75.66 | 82.62 | | *66.00* | 82.00 | | 68.00 | *81.00* |
| P1C1 | *62.90* | 69.46 | 75.99 | 82.37 | *62.00* | 69.00 | 82.50 | *67.00* | 79.00 |
| P1C10 | *68.32* | 76.84 | 82.55 | | *67.00* | 82.00 | | *72.00* | |
| P1E11 | *63.77* | 69.50 | 77.46 | 82.57 | *63.00* | 70.00 | 80.00 | *69.00* | 79.00 |
| P1F2 | *67.75* | 69.81 | 79.76 | 83.24 | *68.00* | 83.00 | | 68.00 | *75.00* |
| P1F8 | *71.76* | *75.80* | 82.71 | | *73.00* | 83.00 | | 68.00 | *84.00* |
| P1G12 | *68.99* | 76.42 | 82.38 | | *68.00* | 82.00 | | *69.00* | |
| P2D10 | *70.57* | 77.40 | 82.44 | | *70.00* | 81.00 | | 68.00 | *79.00* |

*Main Peak*

DSC - Differential Scanning Calorimetry; DSF - Differential Scanning Fluorimetry; DSF* - mAb-carfentanil complex

**Table S2.** DSC and DSF measurements of selected monoclonal antibodies.

| Data collection | Apo C10-S66K Fab | C10-S66K Fab:fentanyl | C10-S66K Fab:carfentanil |
|---|---|---|---|
| Beamline | APS-23IDD | SSRL12-2 | SSRL12-2 |
| Wavelength (Å) | 1.03321 | 0.97946 | 0.97946 |
| Space group | $P6_2$ | $P2_1$ | $P2_1$ |
| Unit cell parameters (Å, °) | a=92.90, b=92.90, c=123.94 | a=71.29, b=73.35, c=95.20 | a= 71.05, b= 73.49, c= 95.09 |
| | $\alpha$=90, $\beta$=90, $\gamma$=120 | $\alpha$=90, $\beta$=90.5, $\gamma$=90 | $\alpha$=90, $\beta$=90.4, $\gamma$=90 |
| Resolution (Å) | 50.00-2.80 (2.85-2.80)[a] | 50.00-1.80 (1.83-1.80)[a] | 95.09 - 1.78 (1.81 - 1.78)[a] |
| Unique Reflections | 14,553 (722)[a] | 87,845 (4,175)[a] | 91,341 (4,542)[a] |
| Redundancy | 7.1 (7.1)[a] | 3.2 (2.8)[a] | 3.2 (3.3)[a] |
| Completeness (%) | 99.5 (96.2)[a] | 97.5 (98.1)[a] | 97.8 (98.7)[a] |
| $<I/\sigma_I>$ | 11.2 (1.1)[a] | 21.1 (3.7)[a] | 28.9 (1.0)[a] |
| $R_{sym}$[b] (%) | 13.9 (142.4)[a] | 7.8 (35.9)[a] | 14.6 (27.3)[a] |
| $R_{pim}$[b] (%) | 5.6 (56.8)[a] | 5.2 (24.9)[a] | 9.7 (17.6)[a] |
| $CC_{1/2}$[c] (%) | 81.0 (29.8)[a] | 94.6 (77.6)[a] | 96.5 (86.2)[a] |
| **Refinement statistics** | | | |
| Resolution (Å) | 49.10-2.82 | 34.72-1.80 | 44.72-1.78 |
| Reflections (work) | 14,525 | 87,773 | 91,258 |
| Reflections (test) | 758 | 4,487 | 2,001 |
| $R_{cryst}$[d] / $R_{free}$[e] (%) | 25.7/30.5 | 17.8/21.6 | 17.8/20.7 |
| **No. of atoms** | | | |
| Fab | 3082 | 6437 | 6470 |
| Drug | - | 106 | 118 |
| Water | - | 773 | 806 |
| Protein G | 410 | - | - |
| **Average B-value (Å$^2$)** | | | |
| Fab | 81 | 20 | 20 |
| Drug | - | 16 | 26 |
| Water | - | 29 | 29 |
| Protein G | 102 | - | - |
| Wilson B-value | 72 | 16 | 16 |
| **RMSD from ideal geometry** | | | |
| Bond length (Å) | 0.006 | 0.014 | 0.007 |
| Bond angle (°) | 1.14 | 1.55 | 0.98 |
| **Ramachandran statistics**[f] | | | |
| Favored (%) | 90.34 | 97.63 | 97.75 |
| Outliers (%) | 1.72 | 0.00 | 0.00 |

a. Numbers in parentheses refer to the highest resolution shell.
b. $R_{sym} = \Sigma_{hkl} \Sigma_i | I_{hkl,i} - <I_{hkl}> | / \Sigma_{hkl} \Sigma_i I_{hkl,i}$ and $R_{pim} = \Sigma_{hkl} (1/(n-1))^{1/2} \Sigma_i | I_{hkl,i} - <I_{hkl}> | / \Sigma_{hkl} \Sigma_i I_{hkl,i}$, where $I_{hkl,i}$ is the scaled intensity of the i[th] measurement of reflection h, k, l, $<I_{hkl}>$ is the average intensity for that reflection, and *n* is the redundancy.
c. $CC_{1/2}$ = Pearson correlation coefficient between two random half datasets.
d. $R_{cryst} = \Sigma_{hkl} | F_o - F_c | / \Sigma_{hkl} | F_o |$ x 100, where $F_o$ and $F_c$ are the observed and calculated structure factors, respectively.
e. $R_{free}$ was calculated as for $R_{cryst}$, but on a test set comprising 5% of the data excluded from refinement.
f. From MolProbity[15].

**Table S3.** X-ray data collection and refinement statistics.

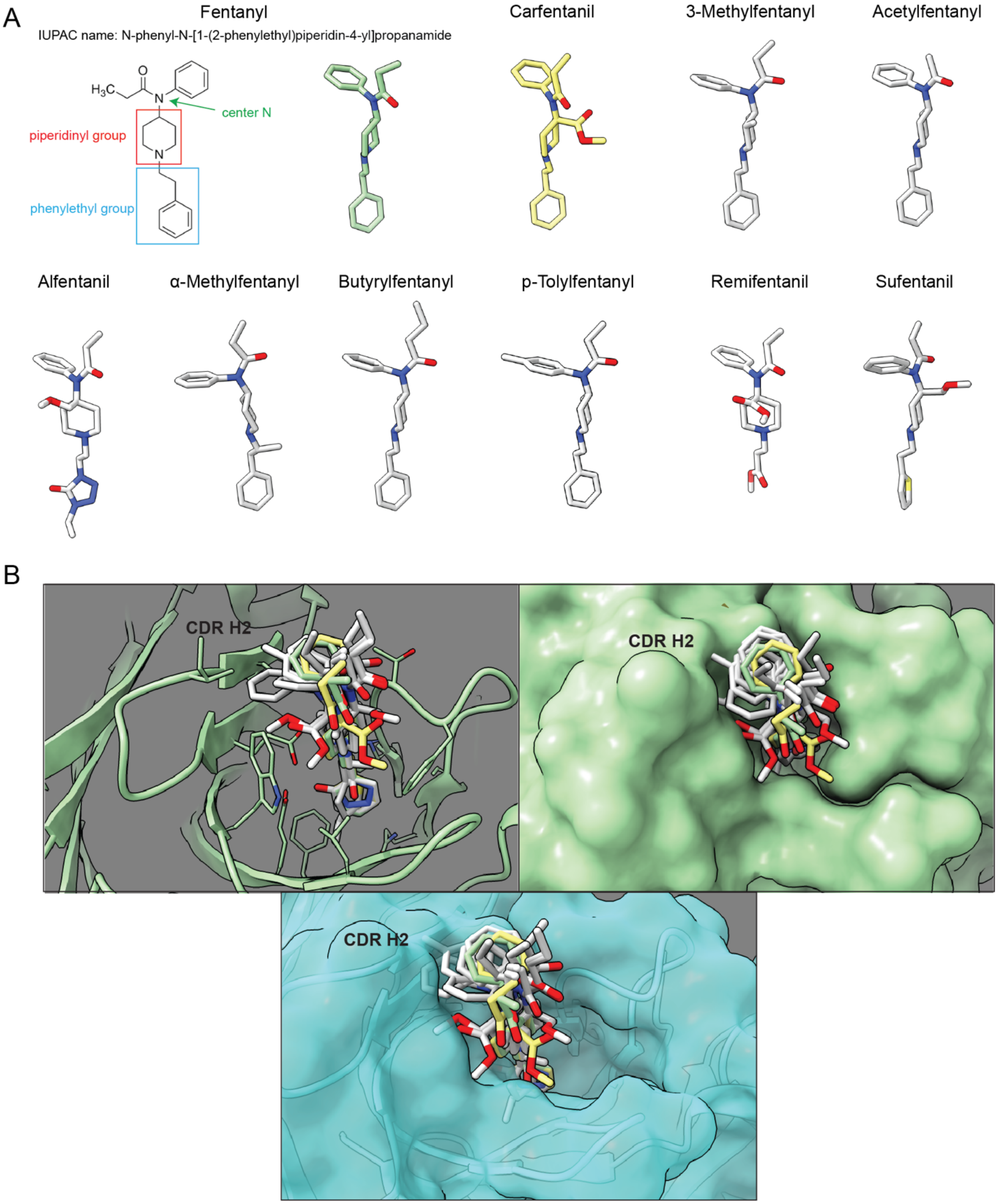


**Supplemental Figure S2.** Fentanyl analogs and the possibility of their interactions with C10-S66K. (A) Structures of fentanyl and fentanyl analogs represented as sticks (all structures were obtained from PubChem). (B) Top-left, alignment of all drugs to fentanyl in the C10-S66K:fentanyl structure. The antibody is shown as cartoons with side chains of the paratopes as sticks. Top right, the same alignment with C10-S66K shown as a green surface. Bottom, the same alignment of the drugs but overlaid on the apo C10-S66K structure, shown as cyan cartoons with a transparent cyan surface.

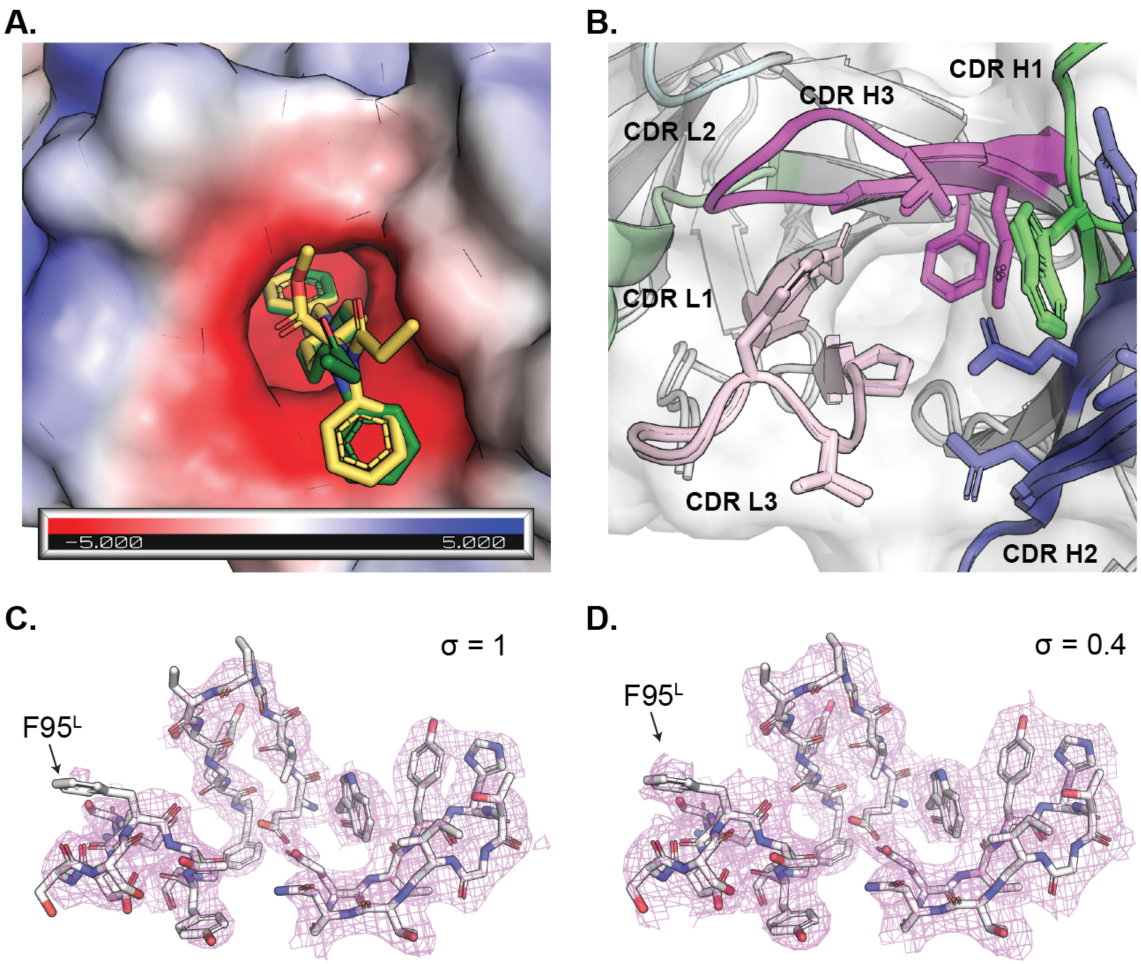


**Supplemental Figure S3.** Electrostatic surface, CDRs, and electron density from C10-S66K paratope. (A) Alignment of stick representations of fentanyl (green) and carfentanil (yellow) from the antibody-bound complexes displayed with the surface of C10-S66K from the fentanyl-bound complex colored by electrostatics (blue as positive charge and red as negative, estimated in PyMOL). (B) Alignment of the cartoon and surface representations of the C10-S66K binding sites for fentanyl and carfentanil. Complementarity-determining regions (CDRs) are colored as shown and surfaces are shown in light grey. (C) Electron density of the paratope of C10-S66K in the apo structure. The apo C10-S66K structure is shown as cartoons with side chains of residues in the paratope as sticks. The $2F_O$-$F_C$ map of C10-S66K at a contour level ($\sigma$) of 1 is shown as magenta mesh. (D) The same representation as in (C) but with the $2F_O$-$F_C$ map at $\sigma = 0.4$. The side chain of $F95^L$ is highlighted with the black arrow.

| scFv mAb | Tm1 (°C) | Tagg 266 (°C) | Z Avg Dia (nm) | PDI |
|---|---|---|---|---|
| wild-type P1A4 | 31.4 | 42.9 | 5.56 | 0.67 |
| C10-S66K | 54.1 | 47.4 | 4.99 | 0.84 |

**Table S4.** Biophysical profile of scFv antibodies using the Uncle System.

| Synthetic Opioid | $k_a$ (1/(Ms) | $k_d$ (1/s) | $K_D$ (M) |
|---|---|---|---|
| Carfentanil | 7.144E+06 | 4.929E-04 | 6.899E-11 |
| Fentanyl | 1.064E+07 | 2.423E-03 | 2.277E-10 |
| Acetylfentanyl | 1.291E+07 | 2.457E-03 | 1.903E-10 |
| Butyrylfentanyl | 2.661E+07 | 3.519E-03 | 1.323E-10 |
| p-Tolylfentanyl | 6.896E+06 | 1.229E-03 | 1.781E-10 |
| 3-Methylfentanyl | 1.083E+06 | 9.536E-03 | 8.809E-09 |
| α-Methylfentanyl | 8.144E+06 | 5.784E-03 | 7.102E-10 |
| Sufentanil | 3.034E+06 | 5.792E-02 | 1.909E-08 |

**Table S5.** Binding parameters of C10-S66K monoclonal antibody.

| mAb | $t_{1/2}$ (h) | AUC (μg/mL*h) | $C_{max}$ (μg/mL) |
|---|---|---|---|
| C10-S66K scFv | 0.88 | 13 | 7.4 |
| C10-S66K scFv-ABD | 46.5 | 1582 | 31.9 |

**Table S6.** Calculated pharmacokinetic parameters for C10-S66K scFv and C10-S66K scFv-ABD.
